# Llgl1 mediates timely epicardial emergence and establishment of an apical laminin sheath around the trabeculating cardiac ventricle

**DOI:** 10.1101/2023.08.14.553249

**Authors:** Eric JG Pollitt, Christopher J Derrick, Juliana Sánchez-Posada, Emily S Noël

## Abstract

During heart development, the embryonic ventricle becomes enveloped by the epicardium, a layer of mesothelium which adheres to the outer apical surface of the heart. This is concomitant with onset of ventricular trabeculation, where a subset of cardiomyocytes lose apicobasal polarity and delaminate basally from the ventricular wall, projecting into the cardiac lumen to begin building the muscle mass necessary for adult cardiac function. Lethal(2) giant larvae homolog 1 (Llgl1) regulates the formation of apical cell junctions and apicobasal polarity, and we investigated its role in ventricular wall maturation, including trabeculation and epicardial establishment. We found that *llgl1* mutant zebrafish embryos exhibit aberrantly positioned cardiomyocytes during early trabeculation, some of which extrude apically into the pericardial space. While investigating apical cardiomyocyte extrusion we identified a basal to apical shift in laminin deposition in the ventricular wall. Initially laminin deposition occurs on the luminal (basal) surface of the heart but concomitant with the onset of trabeculation basal laminin is removed and is instead deposited on the exterior (apical) surface of the ventricle. We find that epicardial cells express several laminin subunits as they adhere to the ventricular wall, and show that the epicardium is required for laminin deposition on the ventricular surface. In *llgl1* mutants the timing of the basal-apical laminin shift is delayed, in line with a delay in establishment of the epicardial layer. Analysis of earlier epicardial development reveals that while both Llgl1 and laminin are not required for specification of the proepicardial organ, they are instead required for dissemination of epicardial cells to the ventricular surface. Together our analyses reveal an unexpected role for Llgl1 in correct timing of epicardial development, supporting integrity of the myocardial wall during early trabeculation.

## Introduction

During cardiac development the ventricular wall is initially two cell layers thick, comprising an outer layer of cardiomyocytes (CMs), and an inner layer of endocardium. As the ventricular wall matures trabeculation is initiated, a process which is crucial for building muscle mass and improving pumping efficiency in the heart. During trabeculation cardiomyocytes delaminate from the basal side of the myocardial wall into the lumen of the chamber, forming trabecular seeds in the lumen of the developing ventricle, from which trabecular ridges subsequently elaborate (Gunawan et al., 2021; Staudt et al., 2014). Onset of trabeculation occurs at around 55-60hpf (hours post fertilisation) in zebrafish, around the same stage as the development of the epicardium, a conserved mesothelial layer which envelops the outer ventricular surface and contributes multiple cell types to the mature heart including coronary cells and fibroblasts (Cao et al., 2020; Peralta et al., 2014). Epicardial development is initiated by formation of the proepicardium through delamination of cells in the dorsal pericardium, forming proepicardial clusters at the venous pole and atrioventricular canal. Epicardial cells are then released from the proepicardial organ, move through the pericardial cavity, and subsequently attach to the myocardium. The concurrent timing of epicardial establishment with trabeculation suggests there may be links between these two processes. While previous studies have demonstrated that loss of epicardium doesn’t impact on early trabeculation (Boezio et al., 2023), zebrafish with mutations in genes required for epicardial development do exhibit defects in ventricular wall integrity, characterised by aberrant apical extrusion of cardiomyocytes into the pericardial space.

The ventricular myocardial wall comprises an epithelium with apicobasal polarity. The basal surface of ventricular CMs is the interior, luminal, ventricular wall (adjacent to the endocardial layer and ventricular lumen), from which cardiomyocytes will delaminate to form trabecular seeds (Jiménez-Amilburu et al., 2016). Conversely, the apical CM surface is the external, abluminal ventricular wall, facing the pericardial cavity (to which the epicardium will adhere). Establishment and maintenance of apicobasal polarity in epithelial cells is also regulated by interactions between three complexes: the Crumbs, Scribble, and Par complexes (Martin et al., 2021). The onset of trabeculation is preceded by the relocalisation of the apical protein Crumbs 2a (Crb2a) from apical CM junctions to the apical surface of CMs, suggesting trabeculation is accompanied by destabilisation of apical cell-cell junctions, which facilitates basal delamination (Jiménez-Amilburu and Stainier, 2019). *crb2a* mutant embryos display multilayered ventricular wall cardiomyocytes that fail to undergo trabeculation, demonstrating that classic regulators of apicobasal polarity are important in ventricular wall maturation, however the role of the other apicobasal complex proteins in ventricular wall polarity and maturation remains to be determined. Apicobasal polarity of epithelia is also often typically supported by the basal deposition of the ECM component laminin (Buckley and St Johnston, 2022; Matlin et al., 2017), and therefore it is assumed that a laminin-rich basement membrane is present on the luminal surface of the heart, although this has not been directly shown during cardiac development prior to trabeculation. However, basal delamination and extrusion of cells in different biological contexts (e.g. cancer) is accompanied by degradation of basal laminin, and basement membrane ECM in general (Akhavan et al., 2012; Banerjee et al., 2022), suggesting that if laminin is basally deposited in the early ventricle, this may need to be (locally) degraded to support trabecular seeding. Consequently, there are key questions to be addressed around whether cardiomyocyte delamination requires remodelling of basal laminin, how cell delamination remains directional and whether regulation of apicobasal polarity links these phenomena. In Drosophila, Lgl (Lethal giant larvae) forms part of the basolateral Scribble complex and regulates timely redistribution of the apical Crumbs complex in epithelia during larval development (Bonello and Peifer, 2019; Martin et al., 2021), suggesting that zebrafish homologs of *Lgl* may also be important for apicobasal polarity in the ventricular wall. Zebrafish have two *Lgl* homologues, *llgl1* and *llgl2*, and *llgl1* has previously been shown to be required for early stages of heart morphogenesis (Flinn et al., 2020). However, whether *llgl1* plays a role in ventricular wall development has not been examined.

In this study we describe requirements for *llgl1* in maintaining ventricular wall integrity at the onset of trabeculation, and in facilitating timely establishment of the epicardial layer around the ventricle. We reveal that the epicardium deposits a layer of laminin at the apical CM surface of the ventricle, and provide evidence for a similar requirement for laminin in epicardial development and maintenance of ventricular wall integrity. Together our results reveal novel roles for apicobasal regulators in ventricular wall maturation during heart development.

## Results

### Llgl1 regulates ventricular wall integrity and trabeculation

Establishment, maintenance and regulation of apicobasal polarity is important for ventricular wall maturation (Gunawan et al., 2021; Jiménez-Amilburu et al., 2016). *llgl1 (lethal(2) giant larvae homolog 1)* is a homologue of drosophila *Lgl*, a conserved mediator of apicobasal polarity (Martin et al., 2021), and we hypothesised that *llgl1* may also play a role in ventricular wall maturation. To investigate this we generated a novel *llgl1* mutant using CRISPR-Cas9-mediated genome editing, recovering a mutant with a 32bp deletion in exon 2, resulting in a truncated protein lacking all functional domains of Llgl1 (Fig S1A-B). *llgl1* mutants exhibit mild cardiac oedema at 72hpf which resolves in most embryos by 5dpf (Fig S1C-H), and are adult viable. Analysis of heart looping in fixed embryos by mRNA *in situ* hybridisation analysis of *myl7* expression revealed a variable reduction in looping morphogenesis in *llgl1* mutants compared to wild-type from 40hpf onwards (Fig S1I-R), in line with previously-described heart phenotypes in *llgl1* mutants and morphants (Flinn et al., 2020). We further analysed heart morphology using live lightsheet microscopy of *Tg(myl7:LifeActGFP);Tg(fli1a:AC-TagRFP)* double transgenic wild-type and *llgl1* mutant embryos between 55hpf and 120hpf when looping morphogenesis is complete, revealing that *llgl1* mutants continue to exhibit defects in heart morphogenesis (Fig S1S-X).

Analysis of the ventricular wall in *llgl1* mutants after initiation of trabeculation at 120hpf revealed phenotypes ranging from similar trabecular organisation as wild-type (Fig 1B) through to complete disorganisation of the trabeculae, including ventricular CM multilayering (Fig 1C-E). We occasionally observed embryos in which CMs were extruding apically into the pericardial cavity (Fig 1E), suggesting defects in polarity or integrity of the ventricular wall. Apically-extruding cells have been previously described to occur at earlier developmental time points in zebrafish embryos harbouring mutations in the EMT-related gene *snai1b* (Gentile et al., 2021), or in mutants with defects in trafficking of subapical junctional proteins (Grassini et al., 2019). We therefore hypothesised that since Llgl1 is involved in apicobasal polarity it may play a similar role in maintaining integrity of the ventricular wall at earlier developmental stages and analysed the ventricular wall prior to trabeculation at 55hpf and during early trabeculation at 80hpf. At all stages examined we consistently observed CMs that extrude from the apical surface of the ventricle into the pericardial cavity in *llgl1* mutants (Fig 1F-K). At 55hpf we found no significant differences in cell extrusion in *llgl1* mutants, however at 80hpf we observed a significant increase in the number of extruding CMs in *llgl1* mutants compared to wild-type siblings. By 120hpf, when trabeculation is now driven by elaboration of trabecular seeds rather than CM delamination (Samsa et al., 2013), extruding cell number in *llgl1* mutants was comparable to wild-type (Fig 1K). The small number of extruding CMs previously observed in wild-type embryos are primarily located around the ventricular apex proximal to the atrioventricular canal (Gentile et al., 2021), consistent with our analysis of extruding cell location in wild-type siblings at 80hpf (Fig 1L). However, extruding cells in *llgl1* mutants are distributed more broadly throughout the ventricle, including elevated numbers in the outer curvature and ventricular apex, as well as some cells in the inner curvature and outflow tract. Together this suggests that *llgl1* is required for ventricular wall maturation.

**Figure 1.**
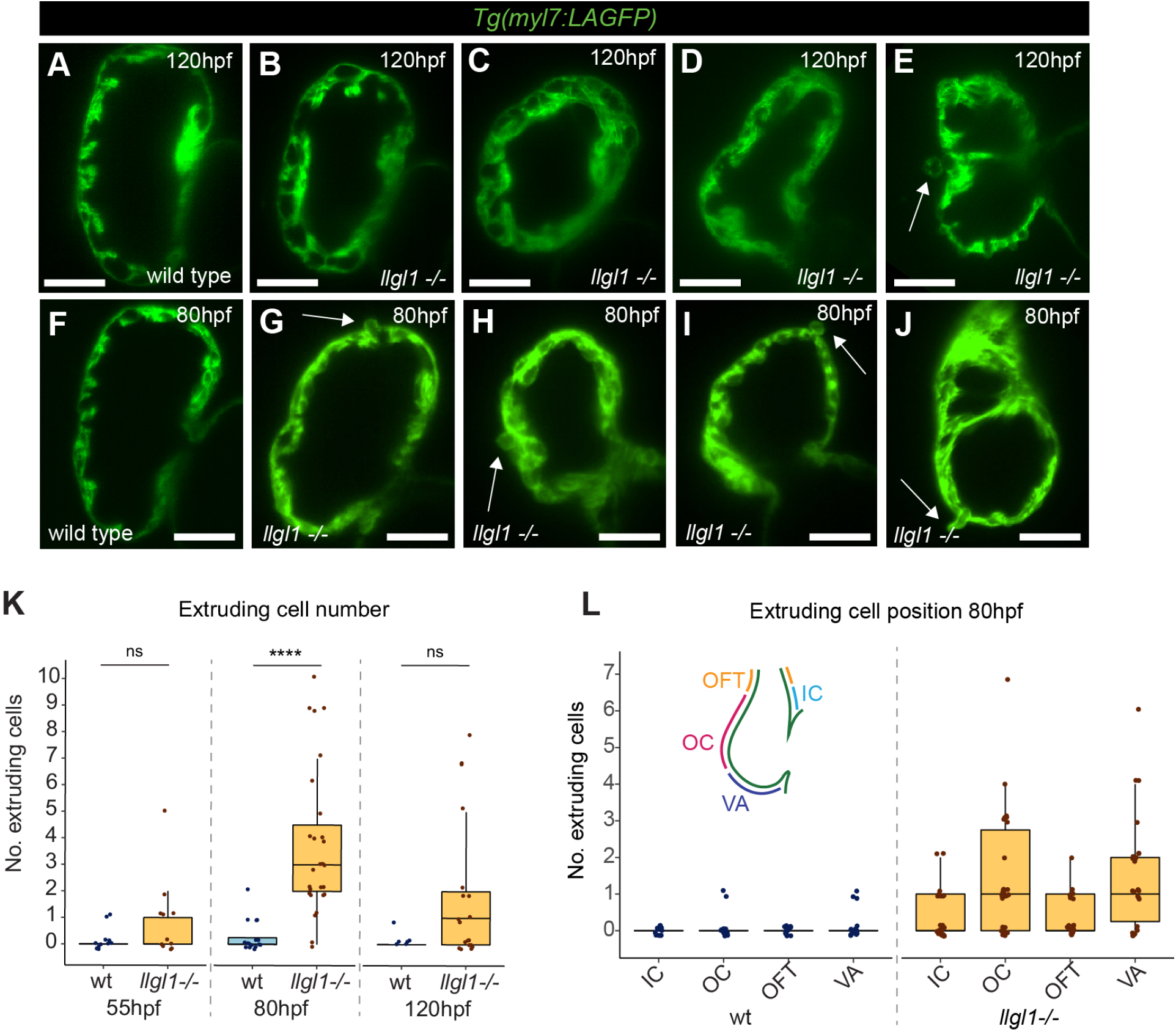
Llgl1 promotes organised trabeculation and ventricular wall integrity. A-J: Live lightsheet z-slices through the ventricle of *Tg(myl7:LifeAct-GFP)* transgenic wild-type and *llgl1* mutant embryos visualising the myocardium at 120hpf (A-E) and 80hpf (F-J). Scale bars: 50μm. Wild-type embryos have regular trabeculae emerging predominantly from the outer curvature of the ventricular wall at 120hpf (A), while *llgl1* mutant embryos exhibit disorganised trabeculae, ranging from wild-type-like (B), through to irregular emergence of trabecular CMs and multilayering of CMs (C-E). Some *llgl1* mutants at 120hpf exhibit apically extruding CMs (arrow, E). At 80hpf in *llgl1* mutants, CMs can be seen aberrantly extruding apically from multiple locations in the ventricular wall (arrows G-J). K: Quantification of extruding cell number in wild-type siblings and *llgl1* mutant embryos at 55hpf (wt, n=11; *llgl1 -/-*, n=11), 80hpf (wt, n=15; *llgl1 -/-*, n= 27) and 120hpf (wt, n=6; *llgl1 -/-*, n=19). L: Distribution of extruding cells in wild-type and *llgl1* mutant embryos at 80hpf. Schematic depicts the location of the outer curvature (OC), ventricular apex (VA), outflow tract (OFT) and inner curvature (IC) in the zebrafish ventricle.

### Loss of llgl1 results in temporal defects in Crumbs redistribution

The ventricular wall exhibits apicobasal polarity prior to and during trabeculation (Jiménez-Amilburu et al., 2016; Jiménez-Amilburu and Stainier, 2019). Loss of the apicobasal polarity regulator Crb2a results in multi layering of polarised CMs, but not apical CM extrusion (Jiménez-Amilburu and Stainier, 2019), but conversely improper CM trafficking of N-cadherin is associated with apical CM extrusion (Grassini et al., 2019). As the *llgl1* mutant displays both ventricular CM multilayering and extrusion, our data suggests a complex relationship between ventricular wall apicobasal polarity and organised CM delamination. Drosophila Lgl is required for timely apical localisation of Crumbs in the larval epithelium (Tanentzapf and Tepass, 2003). This raised the possibility that zebrafish Llgl1 may also be engaged in Crb2a (a Crumbs homolog) redistribution from apical CM junctions to the apical CM membrane during trabeculation (Jiménez-Amilburu and Stainier, 2019). We therefore investigated whether apicobasal polarity in the ventricular wall is disrupted in *llgl1* mutants through quantification of Crb2a distribution across the apical CM surface (Fig 2A-E). In wild-type embryos we identified low levels of Crb2a along the apical CM membrane at 72hpf, with slightly higher junctional than apical Crb2a (Fig 2F, H). Distinctly, *llgl1* mutants have a very strong retention of Crb2a at CM junctions compared to wild-type junctions, or compared to the apical membrane of mutant CMs (Fig 2G,H). However by 80hpf the junctional:apical membrane distribution of Crb2a in *llgl1* mutants has become more comparable with that observed in wild-type CMs, and in general *llgl1* mutants have a slight decrease in overall Crb2a levels (Fig 2H). Together this supports a role for Llgl1 in timely Crb2a protein relocalisation or deposition in ventricular CMs (Fig 2I), although it is unclear whether this Crb2a redistribution is required to maintain CM wall integrity.

**Figure 2.**
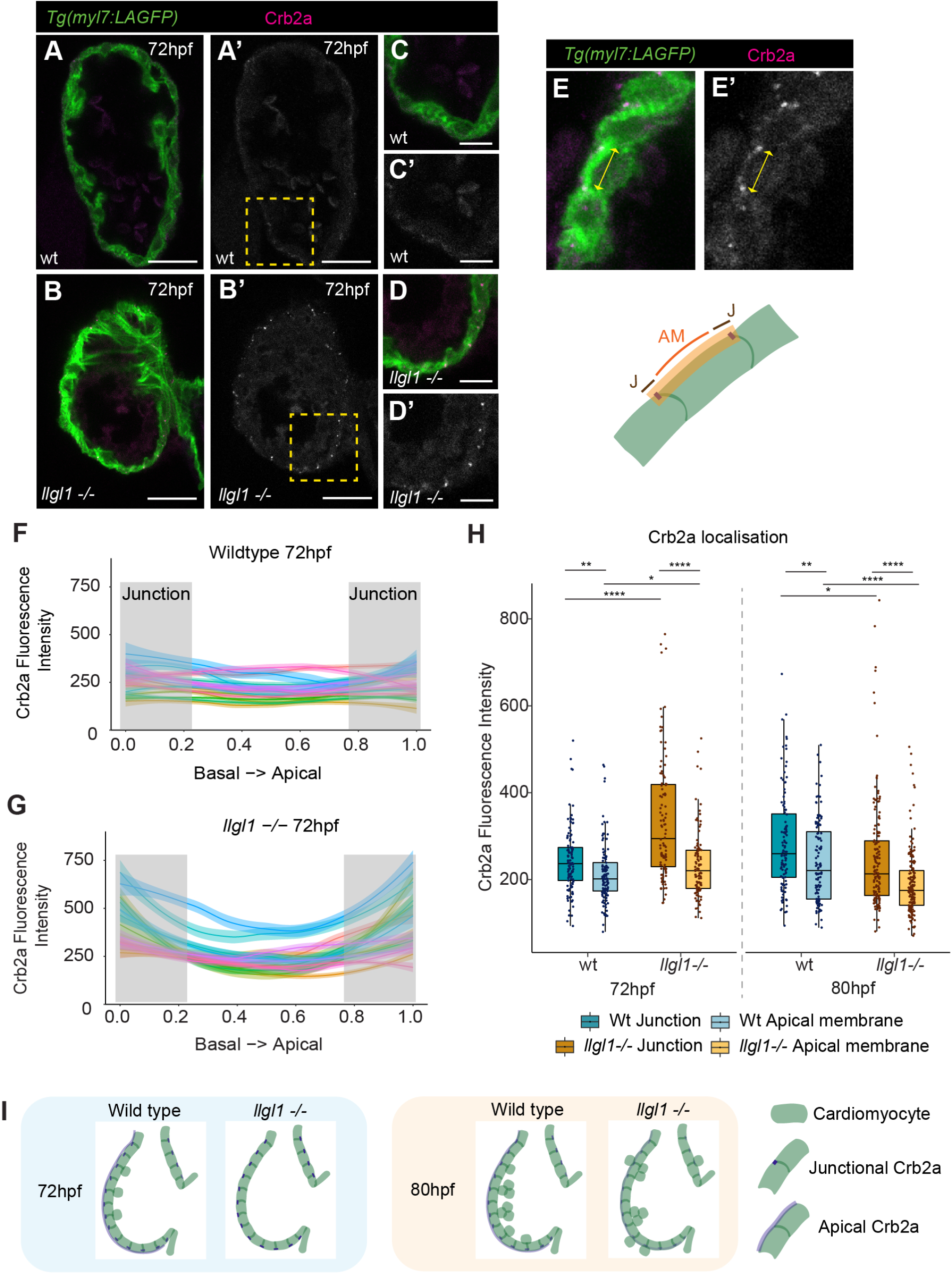
Llgl1 is required for timely apical relocalisation of Crb2a. A-D: Confocal z-slices of the ventricle of *Tg(myl7:LifeAct-GFP)* transgenic embryos visualising the myocardium (green) stained with an anti-Crb2a antibody (magenta). A’-D’ shows Crb2a staining alone. Scale bar = 25µm. C-D show higher magnification of region depicted by yellow boxes in A,B. Scale bar = 10µm. In wild-type embryos at 72hpf low levels of Crb2a are distributed across the apical myocardial membrane (A, C), while in *llgl1* mutants bright Crb2a puncta are observed at apical CM junctions (B, D). E: Example images and schematic showing quantification of Crb2a across the apical CM surface. AM -apical membrane, J - junction. F-H: Quantification of Crb2a intensity across the standardised apical membrane of individual cardiomyocytes in wild-type (F) and *llgl1* mutants (G) at 72hpf. Crb2 appears to be more elevated at the cell boundaries in *llgl1* mutants (G) when compared to wild-type (F). Grey boxes indicate apicobasal positions used to bin data into junctional domains. H: Quantification of junctional/apical Crb2a intensity in wild-type siblings and *llgl1* mutants at 72hpf (wt n=117, *llgl1*-/- n=106) and 80hpf (wt n=112, *llgl1*-/- n=159). At 72hpf wild-type embryos have slightly elevated Crb2a at CM junctions compared to across the apical membrane, but *llgl1* mutants have significantly more junctional Crb2a. By 80hpf overall levels of Crb2a in *llgl1* mutants is reduced compared to wild-types. T-test adjusted for multiple comparisons, **** p<0.0001, *** p<0.001, ** p<0.01, * p<0.05, ns = non significant. I: Schematic depicting dynamics of Crb2a distribution in wild type and *llgl1* mutants.

### Timely establishment of a laminin sheath around the apical ventricular surface requires Llgl1

As well as intracellular polarity complexes, there is a wealth of evidence supporting a role for the extracellular matrix (ECM) in epithelial polarisation, in particular Laminin, a major constituent of the basement membrane (Matlin et al., 2017). Studies have associated breakdown of basement membranes with cell delamination in EMT (Zeisberg and Neilson, 2009) including developmental processes such as neural crest migration (Hutchins and Bronner, 2019). It has been suggested that trabeculation is an EMT-like process (Jiménez-Amilburu et al., 2016; Staudt et al., 2014), and mutations in the EMT regulator *snai1b* results in reduced trabeculation, along with aberrant apical ventricular CM extrusion, similar to that seen in *llgl1* mutants. Together this raised questions around the nature of basement membrane dynamics during trabeculation, the relationship between polarity and apicobasal delamination of ventricular CMs, and the role of *llgl1* in regulating or coordinating these processes. To investigate basement membrane organisation, we analysed laminin deposition in zebrafish hearts prior to and after the onset of trabeculation. At 55hpf, before trabecular seeding is initiated, we found laminin deposition at the luminal basal surface of ventricular CMs (Fig 3A-B’). Once trabecular seeding is underway at 84hpf, we observed a loss of basal laminin in ventricular CMs, however this was unexpectedly accompanied by the deposition of laminin on the outer, apical surface of the heart (Fig 3C-D’). We quantified the dynamics of basal laminin degradation and apical laminin establishment across the apicobasal axis of ventricular CMs (Fig 3E-H), confirming that at 55hpf that laminin is only found on the basal CM surface, whereas by 84hpf laminin has been deposited at the apical CM surface and basal laminin is degraded (Fig 3H). During the onset of trabeculation the ventricular myocardium is a layer of tissue sandwiched between two layers of ECM, therefore we continue to refer to laminin as apical or basal in relation to the myocardium.

**Figure 3.**
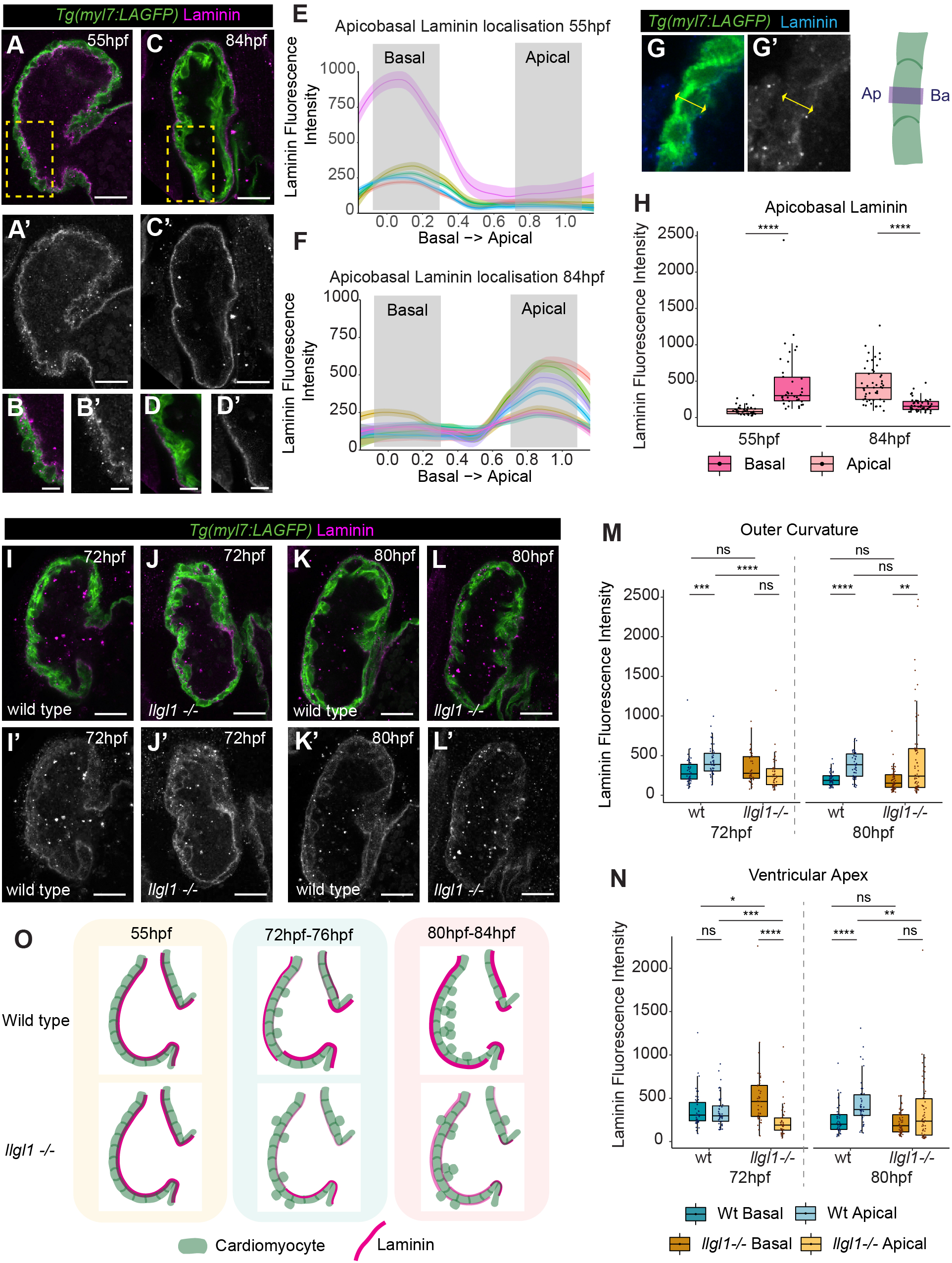
Llgl1 promotes timely deposition of laminin to the apical ventricular surface during early trabeculation. A-D: Confocal z-slices of the ventricle of *Tg(myl7:LifeAct-GFP)* transgenic embryos visualising the myocardium (green) stained with an anti-Laminin antibody (magenta). A-D: merge, A’-D’: Laminin. Laminin is deposited on the luminal basal myocardial surface at 55hpf (A,A’), but is enriched on the apical exterior surface of the myocardium at 84hpf (C-D’). Scale bar = 25µm. B,D: Magnified image of yellow boxed area in A,C. Scale bar = 10µm. E-F: Quantification of Laminin intensity across the standardised apicobasal axis of individual cardiomyocytes at 55hpf (E) and 84hpf (F). Grey boxes indicate apicobasal positions used to bin data into apical and basal domains. G: Example images and schematic showing method for quantifying Laminin across the apicobasal CM axis. Ap - apical, Ba - basal. H: Quantification of Laminin fluorescence intensity at apical and basal positions in ventricular CMs at 55hpf and 84hpf. Each point represents an individual cell, 55hpf, n=38; 84hpf, n=48. Individual t-tests adjusted for multiple comparisons. I-L: Confocal z-slices of the ventricle of *Tg(myl7:LifeAct-GFP)* transgenic embryos visualising the myocardium (green) stained with an anti-Laminin antibody (magenta) in wild-type siblings (I, K) and *llgl1* mutant embryos (J, L). I-L: merge, I’-L’: Laminin. *llgl1* mutants still exhibit basal Laminin at 72hpf (J), and are yet to clearly establish apical Laminin at 80hpf (L). Scale bar = 25µm. M: Quantification of basolateral Laminin intensity in the outer curvature shows that laminin is apically enriched in wild-type siblings from 72hpf onwards (72hpf, n=48; 80hpf, n=49) while *llgl1* mutants still exhibit basal laminin at 72hpf (n=40), and only apical enrichment at 80hpf (n=68). N: Quantification of basolateral Laminin intensity in the ventricular apex reveals wild-type siblings don’t have significant degradation of basal laminin and enrichment of apical laminin until 80hpf (72hpf, n=45; 80hpf, n=42), whereas *llgl1* mutants have significant levels of basal laminin at 72hpf (n=40), and do not have significant apical deposition even by 80hpf. K, L: Kruskal-Wallis test. (n=58). **** p<0.0001, *** p<0.001, ** p<0.01, * p<0.05, ns = non significant. O: Schematic depicting dynamics of Laminin distribution in wild type and *llgl1* mutants.

The timing of the basal to apical shift in laminin localisation in ventricular CMs coincides with the stage at which apical CM extrusion is highest in *llgl1* mutants. Together with our finding that *llgl1* mutants had timely defects in redistribution of the polarity protein Crb2a, this led us to hypothesise that *llgl1* may be required for laminin deposition to the apical CM surface and that this apical laminin may be linked with maintaining integrity of the ventricular wall during trabecular seeding. To determine whether Llgl1 is required for timely laminin deposition to the apical ventricular surface, we analysed laminin deposition in *llgl1* mutants at 72hpf and 80hpf (Fig 3I-L’). We quantified apicobasal CM laminin levels, separating the ventricular wall into outer curvature and ventricular apex, as these were the regions in which we observed the highest number of extruding cells (Fig 1). Interestingly, we noticed that in wild-type embryos, laminin is already apically enriched in the outer curvature at 72hpf (Fig 3M), while at the ventricular apex, apical enrichment of laminin doesn’t occur until around 80hpf (Fig 3N), suggesting a spatiotemporal regulation of laminin deposition to the apical CM surface. *llgl1* mutants have significantly less apical laminin than wild-type siblings in both the outer curvature (Fig 3M) and ventricular apex (Fig 3N) at 72hpf, which in the ventricular apex is also accompanied by a delay in degradation of basal laminin. We started to observe apical enrichment of laminin in the outer curvature of *llgl1* mutants only at 80hpf (Fig 3M), and although basal laminin degradation has accelerated in ventricular apex CMs and apical deposition has begun, laminin is not yet apically enriched (Fig 3N). Levels of apical laminin in *llgl1* mutants at 80hpf are highly variable compared to wild-type siblings, with some mutants exhibiting very high levels of apical laminin for example in the outer curvature, while other mutants are yet to deposit any laminin at the apical surface. Together this suggests that while the spatial dynamics of laminin deposition to the apical ventricular surface are maintained in *llgl1* mutants, Llgl1 is required for the timely establishment of this apical laminin sheath (Fig 3O).

### Epicardial cells deposit laminin on the apical ventricular surface

During zebrafish heart development epicardial coverage of the ventricle is established from around 72hpf onwards (Boezio et al., 2023; Peralta et al., 2014), coinciding with the timing of laminin deposition on the ventricular surface. We therefore investigated whether epicardial cells are the source of apical laminin in the ventricle, first through mRNA *in situ* hybridisation expression analysis of the major cardiac laminin subunits *lamb1a* and *lamc1* between 55hpf and 72hpf (Fig 4A-I). We found that at 55hpf both *lamb1a* and *lamc1* are expressed in a few scattered cobblestone-like cells on or adjacent to the myocardium (Fig 4A,D), with coverage increasing until 72hpf when laminin-expressing cells appear to have flattened out into a thin layer covering the ventricle, adjacent to *myl7-*expressing CMs (Fig 4C,F), morphological changes reminiscent of those previously-described in newly-adhering epicardial cells (Dettman et al., 1997; Pérez-Pomares and de la Pompa, 2011). This suggests that epicardial cells could be the source of laminin deposited on the apical surface of ventricular CMs. To confirm this, we injected embryos with an antisense morpholino oligonucleotide (MO) targeting *wilms tumour 1a (wt1a)* (Perner et al., 2007), a transcription factor expressed in the epicardium and required for epicardial development (Andrés-Delgado et al., 2020; Boezio et al., 2023; Moore et al., 1999; Peralta et al., 2013; Serluca, 2008), and assessed the impact on laminin deposition. To confirm that the *wt1a* morpholino prevented epicardial development, we analysed the expression of Caveolin 1a (Cav1a), a structural component of caveolae expressed in epicardial cells (Grivas et al., 2020) in control MO and *wt1a* MO-injected embryos at 80hpf. While control MO-injected embryos have Cav1a-expressing epicardial cells (Fig 4J), *wt1a* morphant embryos have no epicardial cells at 80hpf (Fig 4L), confirming efficacy of the *wt1a* MO. Corroborating our observation that mRNA encoding laminin subunits is expressed in epicardial cells, analysis of apical laminin deposition revealed that *wt1a* morphants lose apical laminin when compared to control injected (Fig 4K,M), confirming that the epicardium is the source of laminin deposited on the ventricular surface.

**Figure 4.**
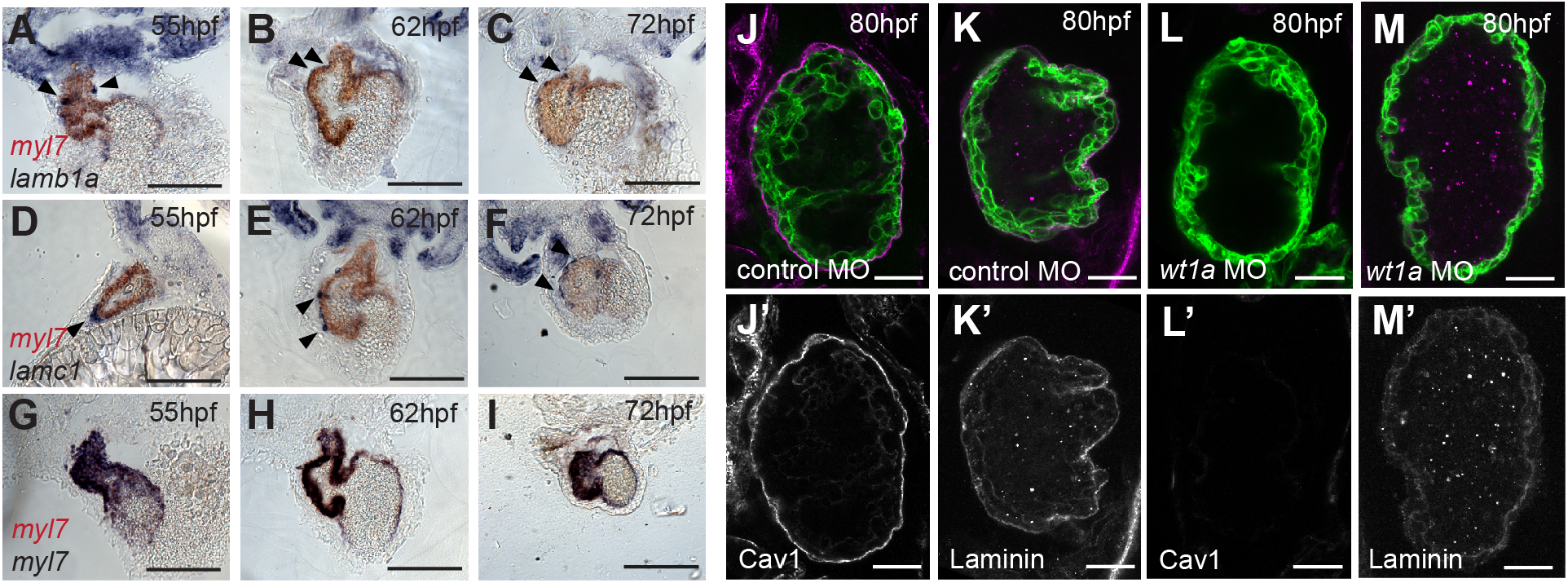
The epicardium deposits laminin onto the apical ventricular surface. A-I: Sections through the hearts of two-colour mRNA *in situ* hybridisation analysis of *lamb1a (*A-C), *lamc1* (D-F), and *myl7* controls (G-I) in blue, in combination with *myl7* expression (red) to highlight the myocardium, between 55hpf and 72hpf (n=3 per gene/stage). Both *lamb1a* and *lamc1* are expressed in epicardial cells adjacent to/on the surface of the myocardium (arrowheads), contrasting with the *myl7* blue/red control *in situ* demonstrating the appearance of co-localising myocardial stains. Scale bars: 200μm. A-C, E-I - coronal sections, D - transverse section. J-M: Confocal z-slices of the ventricle of 80hpf *Tg(myl7:HRAS-GFP)* transgenic embryos injected with either control MO (J, K) or *wt1a* MO (KL M), and stained with either an anti-Cav1a antibody (magenta, J, L) or anti-Laminin antibody (K, M). Scale bar = 25µm. Control MO-injected embryos exhibit expression of both Cav1a (n=9/10) and Laminin (n=10/10) at the apical CM surface (J, K), while *wt1a* MO-injected embryos show loss of both Cav1a (L, n=9/9) and Laminin expression (M, n=10/10).

### Llgl1 is required for timely emergence of the epicardium from the dorsal pericardium

We next investigated whether *llgl1* mutants exhibit defects in epicardial attachment which could account for the delay in apical laminin establishment by analysing the expression of Cav1a in wild-type and *llgl1* mutant embryos. At 76hpf all wild-type siblings exhibited full epicardial coverage of the ventricle (Fig 5A). However, *llgl1* mutants display variability in epicardial coverage, with some mutants having either full (Fig 5C) or almost/completely absent epicardial coverage (Fig 5D) while the majority exhibiting partial epicardial coverage, often with large patches of epicardium missing at the ventricular apex (Fig 5B). However by 96hpf the majority of *llgl1* mutants have full epicardial coverage of the ventricle comparable with wild-type siblings (Fig 5E,F), suggesting that while *llgl1* is not required for epicardial specification, it is instead required for timely dissemination of epicardial cells. To confirm this, we analysed epicardial development at 55hpf when proepicardial clusters are forming from the dorsal pericardium at the venous pole and atrioventricular canal, and beginning to colonise the ventricle (Andrés-Delgado et al., 2019; Boezio et al., 2023; Serluca, 2008). mRNA *in situ* hybridisation analysis of *wt1a* expression revealed that proepicardial cells are positioned in the dorsal pericardium in *llgl1* mutants similar to wild-type siblings (Fig 5G, I). However, while in wild-type siblings epicardial cells had begun to emerge from the proepicardium and adhere to the ventral ventricular surface (Fig 5H), in *llgl1* mutants very few epicardial cells had started colonising the ventricle (Fig 5J, K). Previous studies have shown that cardiac contractility is important for epicardial colonisation of the ventricle (Peralta et al., 2013), however we observe no defects in heart rate in *llgl1* mutants at 72hpf (Fig S2), suggesting the epicardial defect in *llgl1* mutants is independent from cardiac function. Together this supports the hypothesis that while *llgl1* is not required for proepicardial specification, it is required for timely epicardial colonisation of the ventricle and subsequent deposition of laminin onto the apical surface of ventricular CMs (Fig 5L).

**Figure 5.**
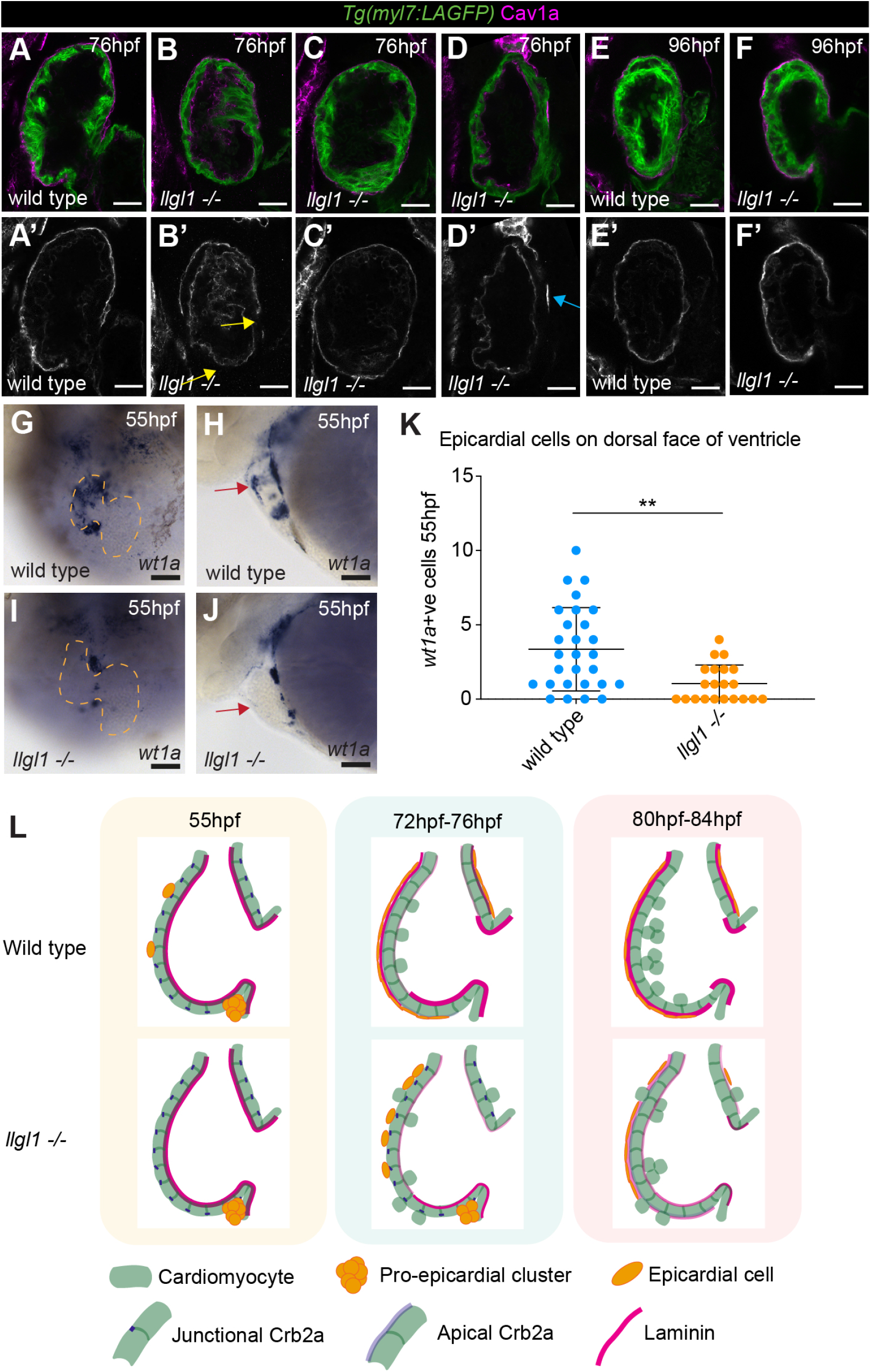
Llgl1 is required for timely epicardial emergence and ventricular colonisation. A-F: Confocal z-slices through the ventricle of *Tg(myl7:LifeAct-GFP)* transgenic embryos visualising the myocardium (green) stained with an anti-Cav1a antibody (magenta) highlighting the epicardium at 76hpf (A-D) and 96hpf (E-F). Wild-type embryos show full epicardial coverage at 76hpf (A, n=12/12), whereas only a small subset of *llgl1* mutants have full epicardial coverage (C, n=3/15) while some have no only a few epicardial cells attached (D, blue arrow, n=3/15) and the majority have partial epicardial coverage (B, yellow arrow, n=9/15). By 96hpf Cav1a-positive epicardium surrounds the ventricle in both wild-type (E, n=4/4), and the majority of *llgl1* mutant embryos (F, n=5/6). Scale bar = 25µm. G-J: mRNA *in situ* hybridisation analysis of *wt1a* expression in wild-type and *llgl1* mutant embryos at 55hpf, G,I - ventral view, dotted line outlines the heart, H,J - lateral view. Scale bar = 50µm. Both wild-type and *llgl1* mutant embryos express *wt1a* in proepicardial cells in the dorsal pericardium, however while wild-type embryos have *wt1a* positive cells attached to the ventral ventricular wall (red arrowhead, H), these cells are reduced or absent in *llgl1* mutants (red arrowhead, J). K: Quantification of the number of epicardial cells attached to the ventral ventricular wall in wild-type (n=28) and *llgl1* mutants (n=21). Comparative analysis performed using T-test, ** p<0.01. L: Schematic model of epicardium, Laminin and Crb2a dynamics between 55hpf and 84hpf.

### Laminin is required for ventricular wall integrity and epicardial development

Previous studies have identified that *wt1a* and *tcf21* mutants, which exhibit epicardial defects, also display aberrant apical extrusion of cardiomyocytes from the ventricular wall, supporting a hypothesis that epicardial coverage of the ventricle maintains myocardial wall integrity during trabecular seeding. However, analysis of hypomorphic alleles in which some epicardial cells remain suggests that the presence of epicardial cells alone is not sufficient to prevent CM extrusion (Boezio et al., 2023). Our observation that the delay in apical laminin deposition in *llgl1* mutant embryos is coincident with the stage at which aberrant ventricular CM extrusion is highest similarly suggests that establishment of an epicardially-derived apical laminin sheath around the outer surface of the ventricular myocardium may help maintain the integrity of the ventricular wall during early trabeculation. Supporting this we analysed whether apically extruding CMs in either wild-type or *llgl1* mutant embryos were associated with less apical laminin. We quantified laminin levels across the apical membrane of both extruding cells and adjacent cells either side of the extruding cell (Fig S3A), and consistent with our hypothesis, we find that extruding cells in wild-type embryos have significantly less laminin than their neighbours (Fig S3B). Despite *llgl1* mutants having lower levels of apical laminin in general, we find the same dynamic as wild-type, where extruding CMs have less laminin than their neighbours. Together this suggests that apical laminin could maintain integrity of the ventricular wall by preventing apical cell extrusion during early trabeculation. To investigate this further we analysed ventricular wall integrity in *lamb1a* mutants, which carry a mutation in a laminin beta 1a subunit (Derrick, Pollitt, et al., 2021), and which we have shown is expressed in epicardial cells (Fig 4). Live lightsheet imaging of the ventricular wall at 76hpf revealed a significant increase in the number of apically extruding ventricular CMs in *lamb1a* mutants when compared to wild-type siblings (Fig S3C-H), albeit at lower frequency than in *llgl1* mutants. Together this suggested that laminin is required to maintain integrity of the ventricular wall.

To confirm that we were impacting apical laminin deposition but not epicardial development we analysed Cav1a expression in *lamb1a* mutants. Surprisingly, and similar to our findings in *llgl1* mutants, the majority of *lamb1a* mutants exhibit partial epicardial coverage at 76hpf when compared with wild-type siblings, with frequent gaps observed in the epicardium (Fig 6A-D), suggesting that laminin is required for epicardial development. Confirming this, we also investigated Cav1a expression in *lamc1* mutants, which carry mutations in the laminin gamma1 subunit (Odenthal et al., 1996; Parsons et al., 2002). *lamc1* mutants exhibit a more severe epicardial phenotype than *lamb1a* mutants, with most mutants having a complete loss of epicardial attachment to the ventricle (Fig 6E,F). This epicardial defects in laminin mutants could result either from a failure in epicardial cells to remain adhered to the ventricular surface at 72hpf if they are unable to produce laminin or that, similar to Llgl1, laminin is also required earlier for epicardial emergence from the proepicardium. To investigate the latter, we analysed *wt1a* expression in *lamb1a* and *lamc1* mutants at 55hpf and found that similar to *llgl1* mutants both laminin mutants exhibit defects in epicardial cell emergence (Fig 6G-N), including retention of proepicardial cells in the dorsal pericardium, and a reduction in epicardial cells positioned on the ventral ventricular surface.

**Figure 6.**
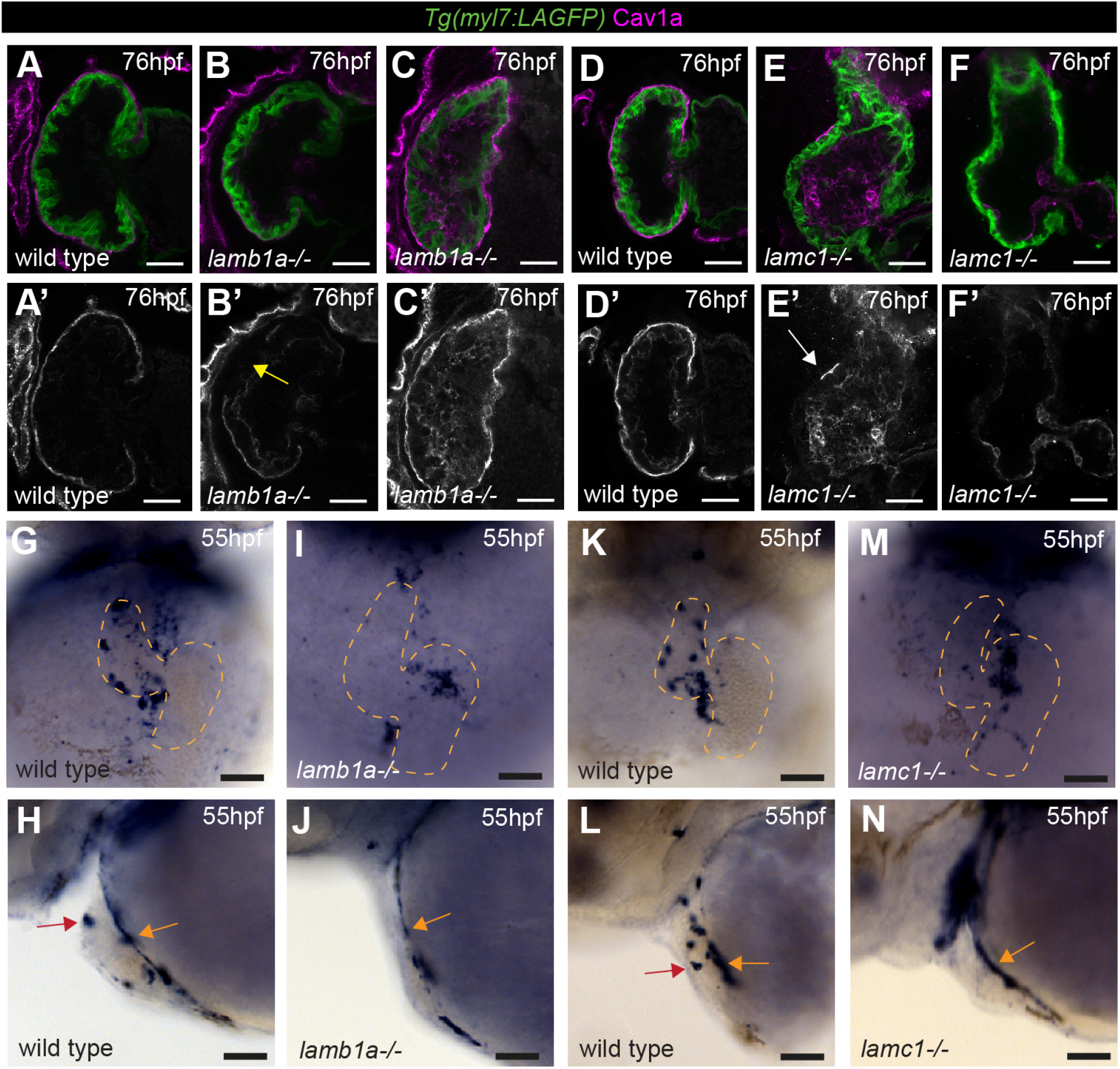
Laminin promotes epicardial emergence and ventricular colonisation. A-F: Confocal z-slices through the ventricle of *Tg(myl7:LifeAct-GFP)* transgenic embryos visualising the myocardium (green) stained with an anti-Cav1a antibody (magenta) highlighting the epicardium at 76hpf in wild-type siblings (A, D), *lamb1a* mutants (B, C), and *lamc1* mutants (E, F). Scale bar = 25µm. wild-type embryos show full epicardial coverage at 76hpf (A, n=12/12, D, n=11/11), while both *lamb1a* and *lamc1* mutants show defects in epicardial coverage. The majority of *lamb1a* mutants have only partial epicardial coverage (B, yellow arrow, n=10/13), with a small number of *lamb1a* mutants exhibiting full epicardial coverage (C, n=3/13). *lamc1* mutants have more profound epicardial defects, with the majority having no epicardial cells attached at 76hpf (F, n=6/9), and a small proportion having only a few epicardial cells attached (E, white arrow, n=3/9). G-J: mRNA *in situ* hybridisation analysis of *wt1a* expression at 55hpf in wild-type embryos (G, H, K, L), *lamb1a* mutants (I, J), and *lamc1* mutants (M, N). G, I, K, M - ventral view, dotted line outlines the heart, H,J, L, N - lateral view. Scale bar = 50µm. Similar to wild-type siblings (H, J, n=10 and n=12 respectively), both laminin mutants have *wt1a* expression in proepicardial cells on the dorsal pericardium (L, N, yellow arrows, n=11 and n=12 respectively). Wild-type embryos also have epicardial cells attached to the ventral ventricular wall (H, J, red arrows) but both *lamb1a* and *lamc1* mutants show little attachment of epicardial cells to the ventral ventricle at 55hpf (L, N).

Together this suggests that while both Llgl1 and laminin aren’t required for proepicardial specification, they are both required for timely epicardial colonisation, representing a more general role for apicobasal polarity in emergence of the epicardium from the dorsal pericardium.

### Myocardial llgl1 is not required to maintain ventricular wall integrity

We have shown that *llgl1* is required for timely deposition of laminin on the apical ventricular CM surface through regulation of epicardial emergence from the dorsal pericardium (Fig 5L). This delayed apical laminin deposition coincides with high numbers of aberrant apically-extruding CMs in the ventricular wall (Fig 1), suggesting that deposition of laminin by epicardial cells may help maintain integrity of the ventricle during trabecular seeding. Supporting this, laminin mutants also exhibit extruding ventricular CMs (Fig S3), however they also exhibit a similar defect in epicardial emergence as observed in *llgl1* mutants, making it difficult to dissect whether these are independent requirements for laminin (Fig 6). Similarly, while zebrafish embryos lacking epicardial cells do exhibit extruding ventricular CMs, it has been suggested that this is not due to the physical presence of the epicardium itself (Boezio et al., 2023). This raises further questions about whether the CM extrusion defects in *llgl1* mutants are secondary to epicardial defects, or represent a role for *llgl1* in cardiomyocytes directly regulating wall integrity. *llgl1* is expressed relatively broadly throughout the embryo (Clark et al., 2012), and scRNA-seq analysis of the developing heart at 48hpf and 72hpf reveal it is expressed at low levels in both myocardial and epicardial cells (Nahia et al., 2023), suggesting it could play a role in both cell types.

It has been hypothesised that increased cell density in the myocardium may lead to elevated apical cell extrusion in embryos lacking epicardium (Boezio et al., 2023), and previous studies found that increased cell density leads to delamination of trabecular CMs during trabecular seeding (Priya et al., 2020). However, analysis of ventricular CM internuclear distance in *llgl1* mutants at both 55hpf and 80hpf reveals no reduction in cell size compared to wild-type siblings (Fig S4), suggesting this is not a CM-intrinsic mechanism underlying CM extrusion. To definitively answer whether *llgl1* is required in the myocardium to prevent aberrant CM extrusion, we generated a *Tg(myl7:llgl1-mCherry)* transgenic line, in which an Llgl1-mCherry fusion protein is expressed only in cardiomyocytes. Analysis of Llgl-mCherry localisation in CMs at 55hpf reveals that it is enriched at the basolateral cell membrane (Fig S5A,B), suggesting correct trafficking and localisation of the fusion protein. To determine whether myocardial Llgl1 can rescue the aberrant cell extrusion in *llgl1* mutant embryos, we generated *llgl1* heterozygous adults carrying the *Tg(myl7:Llgl1-mCherry)* transgene, and outcrossed them to *llgl1*;*Tg(myl7:LifeAct-GFP*) heterozygotes to obtain *llgl1-/-;Tg(myl7:llgl1-mCherry)* embryos. We found no difference in extruding CM number between *llgl1* mutants or *llgl1* mutants with the myocardial *llgl1* rescue transgene (Figure S5C), indicating Llgl1 maintains ventricular wall integrity independent of a function in the myocardium. Supporting this, we find no rescue of epicardial defects in the *llgl1-/-;Tg(myl7:llgl1-mCherry)* (Fig S5D-G). Together this suggests that Llgl1 acts specifically in epicardial cells to maintain ventricular wall integrity during heart development.

## Discussion

We describe for the first time laminin dynamics during the onset of trabeculation. We show that prior to trabeculation laminin is deposited at the basal surface of ventricular cardiomyocytes, but at the onset of trabeculation basal laminin is degraded and an epicardially-derived sheath of laminin is instead deposited on the apical surface of the ventricle. Trabeculation has been described as an EMT-like process (Jiménez-Amilburu et al., 2016), and thus breakdown of laminin at the basal ventricular surface is in line with basement membrane degradation in other EMT events from embryonic development to cancer metastasis (Amack, 2021; Banerjee et al., 2022; Kalluri and Weinberg, 2009; Zeisberg and Neilson, 2009). A previous study in zebrafish has described laminin deposition at the basal surface of cardiomyocytes at 5dpf (Marques et al., 2022), and thus it is possible that basal degradation of laminin is only transient to allow CM delamination associated with trabecular seeding, but once this process is complete basal laminin is re-established. The requirement for establishment of an apical layer of laminin is unclear. Although not as common as basal ECM, apically-deposited ECM has been described in several biological contexts (Li Zheng et al., 2020), where it has been reported to act as external barriers on epithelia, or as internal regulators of tissue morphogenesis. Deposition of specifically laminin to apical tissue surfaces is less common, but has been observed in a few specialised epithelia (Koch et al., 1999; Libby et al., 1996; Sorrosal et al., 2010) suggesting functional roles for apical laminin in specific biological contexts, which may not involve formation of typical basement membrane structures. We considered the possibility of a myocardial origin for apical ventricular laminin, but confirmed expression of laminin subunits in the epicardium during early stages of ventricle colonisation along with an epicardial requirement for apical laminin deposition. In line with this, scRNA-seq analysis at 72hpf identified *lamc1* as a marker of epicardial identity (Nahia et al., 2023), and transcriptomic analysis of *wt1a* mutants identified a downregulation of several ECM components including *lama5* (Boezio et al., 2023). In line with the above we therefore speculate that epicardially-deposited apical laminin around the ventricle could promote ventricular morphogenesis, in line with roles for other apical ECMs, and consistent with the described requirement for the epicardium in ventricular morphogenesis (Boezio et al., 2023). Alternatively, apical laminin could act as a barrier around the external ventricular wall maintaining wall integrity and directionality of CM delamination during early trabeculation. Finally, it is also possible that epicardial deposition of laminin may help anchor the epicardium to the myocardial wall after initial attachment through myocardially-derived VCAM1 (Pae et al., 2008; Sengbusch et al., 2002), and subsequently provide survival and mechanical cues to the developing epicardium and myocardium.

In addition to the roles for Llgl1 and laminin in regulating epicardial emergence, we have also shown that both components help maintain integrity of the ventricular wall, with both *llgl1* and *lamb1a* mutants exhibiting apical extrusion of ventricular CMs. Loss of *Lgl* in the Drosophila imaginal wing epithelia has also been associated with aberrant (basal) extrusion of *Lgl* mutant clones (Froldi et al., 2010), although the cellular environment surrounding the cells is an important mediator of this behaviour which is associated with cell competition. Ventricular CM extrusion has also observed in a variety of different zebrafish mutants, including the EMT transcription factor *snai1b (Gentile et al., 2021),* the atypical myosin *myoVb*, which regulates endosome recycling and N-cadherin trafficking in CMs (Grassini et al., 2019), the flow-dependent transcription factor *klf2 (Rasouli et al., 2018)*, the RA-degrading enzyme *cyp26 (Rydeen and Waxman, 2016),* and epicardial transcription factors *wt1a* and *tcf21* (Boezio et al., 2023). The breadth of pathways implicated in cell extrusion suggest maintenance of ventricular wall integrity during heart morphogenesis and ventricular wall maturation is highly complex. Interestingly, these mutants also present with similar morphological defects to *llgl1* mutants, including dysmorphic ventricles (Flinn et al., 2020) and reduced looping morphogenesis. We observed a temporal delay in Crb2 relocalisation in ventricular CMs of *llgl1* mutants embryos in line with previous studies demonstrating Lgl is required for timely apical localisation of Crumbs in Drosophila epithelia, which is linked to morphogenesis (Tanentzapf and Tepass, 2003). This could suggest that defects in apicobasal polarity in the ventricular wall may affect direction or organisation of cell delamination during trabecular seeding, supporting previous studies which show that complete loss of Crb2a results in CM multilayering in the ventricular wall (Jiménez-Amilburu and Stainier, 2019). However, *crb2a* mutants do not exhibit apical CM extrusion, and unravelling the interactions between Lgl1 and Crb2a in the context of ventricular wall organisation and CM delamination may be challenging since disrupting members of one of the polarity complexes can result in partial compensation from another (Tanentzapf and Tepass, 2003). Nevertheless, we hypothesise that the cell extrusion defects and trabeculation defects in *llgl1* mutants may arise through different mechanisms: the former due to delayed epicardial development, and the latter due to polarity defects in the ventricular wall. We also observed extruding CMs in regions of the ventricle from which CMs do not normally extrude in wild-type embryos. A recent preprint used computational modelling to demonstrate that surface geometry plays a role in biasing cell extrusion events, with extrusion more likely to occur from concave surfaces rather than convex ones (Huang et al., 2022). This raises the interesting possibility that changes in ventricle geometry in *llgl1* mutants may be linked to increased CM extrusion, altered positioning of extruding cells, or disorganised trabecular seeding.

Together, these studies provide plausible evidence for either a CM or epicardial requirement for *llgl1* in preventing CM extrusion. Our investigations of the tissue-autonomous requirement for *llgl1* in maintaining ventricular wall integrity revealed that aberrant CM extrusion in *llgl1* mutants is not due to a myocardial requirement for *llgl1*, and rather it is the delayed epicardial attachment that results in temporal CM extrusion. However, a previous analysis of hypomorphic epicardial mutants suggests the epicardium alone does not physically restrain myocardial cells from extruding apically (Boezio et al., 2023), since there is no significant correlation between the number of epicardial cells on the ventricular surface and extruding CMs, and epicardial cells have been observed on extruding cells. We cannot fully account for these differences. However, exploring further the likelihood of extruding CMs being directly associated with an epicardial cell, together with a more granular understanding of the timing of laminin deposition post epicardial cell adhesion, may help better understand this relationship and these discrepancies.

We have also shown that both Llgl1 and laminin are required for timely emergence of epicardial cells from the dorsal pericardium. This suggests that apicobasal polarity plays a general role in epicardial dissemination, which is in line with a previous study demonstrating a role for the PAR polarity complex member Par3 in proepicardial cyst formation (Hirose et al., 2006). It was proposed that Par3 may act to interpret basal cues from the underlying ECM which helps to polarise proepicardial cells to form cysts, which may explain the similar epicardial defects observed in *llgl1* and *lamb1a/lamc1* mutants.

*LLGL1* lies within the Smith-Magenis microdeletion syndrome (SMS) region on chromosome 17 in humans (Rinaldi et al., 2022). Although the most penetrant symptoms of SMS such as sleep disorders are directly associated with heterozygous loss of *RAI1*, linkage studies suggest that loss of other genes within the deletion region contribute to aspects of the syndrome (Edelman et al., 2007; Girirajan et al., 2006). In particular, 20-40% of SMS patients have heart defects, and a further subset of patients exhibit jaw defects, anterior eye defects and intellectual disability, with large variability in the penetrance of these phenotypes. We speculate *LLGL1* may be of importance in this syndrome since: to-date no candidate gene has been identified that may underlie the cardiac defects in SMS patients (Onesimo et al., 2021); *llgl1* mutant embryos have variable expressivity of phenotypes, particularly regarding heart looping, trabeculation and epicardial development, similar to SMS patients (abnormal heart looping is associated with congenital heart defects (Houyel and Meilhac, 2021)); *llgl1* is expressed in the heart, brain, anterior of the eye and jaw, with previous studies identifying ocular defects in *llgl1* mutants (Clark et al., 2012). Together this raises the interesting possibility that loss of *LLGL1* contributes to cardiac defects in SMS patients.

Together our study reveals for the first time the existence of an epicardially-derived layer of laminin on the apical surface of the ventricle during heart development, providing further evidence that the epicardium helps maintain integrity of the ventricular wall during early trabeculation.

## Acknowledgements

We thank Emma Armitage for critical discussion of the data. Confocal imaging was performed at the Wolfson Light Microscopy Facility using the Nikon A1 microscope and Zeiss Z1 lightsheet microscopes (BBSRC ALERT14 award BB/M012522/1 and BHF Infrastructure Grant IG/15/1/31328). Sectioning and mounting of *in situ* hybridisation samples was performed by Fiona Wright, Histology Hub Facility, Division of Clinical Medicine, UoS. E.P and E.N are supported by a British Heart Foundation Fellowship award FS/16/37/32347.

## Author contributions

Conceptualisation, E.P and E.N; Methodology, E.P, J.S-P, E.N; Investigation, E.P and E.N; Resources, E.P and C.D; Formal Analysis, E.P and E.N; Writing – Original Draft, E.N; Writing – Review & Editing, E.P, C.J.D, J.S.P and E.N; Funding Acquisition, E.N; Supervision, E.N.

## Methods

### Zebrafish Maintenance

The following previously described lines were used: AB; *Tg(myl7:lifeActGFP)* (Reischauer et al., 2014); *Tg(fli1a:AC-TagRFP)^sh511^* (Savage et al., 2019), *Tg(-5.1myl7:DsRed2-NLS)^f2^* (Rottbauer et al., 2002); *lamb1a^sh590^* (Derrick, Pollitt, et al., 2021), *lamc1^sa379^* (Kettleborough et al., 2013). Embryos were maintained in E3 medium (5 mM NaCl, 0.17 mM KCl, 0.33 mM CaCl2, 0.33 mM MgS04) at 28.5°C and were staged as per standard protocols (Kimmel et al., 1995). Embryos older than 24 hpf were transferred into E3 medium containing 0.003% 1-phenyl 2-thiourea (PTU, Sigma-Aldrich P7629). Animal work was approved by the local Animal Welfare and Ethical Review Body (AWERB) at the University of Sheffield, conducted in accordance with UK Home Office Regulations under PPLs 70/8588 and PA1C7120E, and in line with the guidelines from Directive 2010/63/EU of the European Parliament on the protection of animals used for scientific purposes.

### Generation of the llgl1 mutant line

The *llgl1* mutant zebrafish line was generated using a CRISPR guide RNA designed to target Exon 2 of *llgl1* (ENSDART00000003511.11). A CRISPR target sequence (5’- GGCTATTGGAACTAAATCAGGGG-3’) in Exon 2 was identified using CHOPCHOP (Labun et al., 2016; Montague et al., 2014) and the reverse complement inserted into an ultramer scaffold as previously described (Hruscha et al., 2013) for T7 amplification: (5’- AAAGCACCGACTCGGTGCCACTTTTTCAAGTTGATAACGGACTAGCCTTATTTTAACTTG CTATTTCTAGCTCTAAAAC**CTGATTTAGTTCCAATAGCC**CTATAGTGAGTCGTATTACG C-3’). The ultramer was amplified by PCR (F: 5′-GCGTAATACGACTCACTATAG-3′, R: 5′- AAAGCACCGACTCGGTGCCAC-3′) and used as a template for *in vitro* transcription using MEGAshortscript T7 kit (Ambion/Thermo Fisher Scientific AM1354). 2ng gRNA was injected together with 1.9nM Cas9 protein (New England Biolabs M0386T) and 10% Phenol Red (Sigma-Aldrich P0290) into the yolk at the one-cell stage, and injected F0 embryos were raised to adulthood. F0 founders were outcrossed to wild-type, and resulting embryos genotyped by PCR to amplify the region of *llgl1* targeted for mutagenesis (F: 5′- GTCGGGATTGCTCTGAATAGAT -3′, R: 5′- AAGGATACATTTTGATGGCCC -3′), with mutations analysed by Sanger sequencing. A 32bp coding sequence deletion allele was recovered, designated *llgl1^sh598^*, with the deletion region indicated by brackets: 5’- TATGATCCCA[AACTGCAGCTTATGGCTATTGGAACTAAATCA]GGGGCCATCAAAAT-3’. The deletion generates a premature stop codon in exon 2. F0 founders transmitting this mutation was outcrossed to *Tg(myl7:LifeActGFP)* and their offspring raised to adulthood. Phenotypic analyses were carried out on embryos generated from F2 or F3 adults.

### Generation of the Tg(myl7:llgl1-mCherry) transgenic zebrafish line

The *llgl1* full coding sequence (minus the termination codon) was amplified from wild-type cDNA using the following primers: F: 5′-ATGATGAAGTTTAGGTTCAGACGGC-3′; R: 5′- TCAGTTGATGAGGATTCCAGCAGAT-3′. This PCR product was further amplified using primers containing AttB sequences for Gateway cloning at the 5’ end of both primers, and a Kozak sequence before the initiating methionine in the forward primer: F: 5′- GGGGACAAGTTTGTACAAAAAAGCAGGCTTCGCCGCCACCATGATGAAGTTTAGGTTCAGACGGCAGGGAAATGACCCTCATCGT-3′; R: 5′- GGGGACCACTTTGTACAAGAAAGCTGGGTTTGATCCTCCTCCTCCTGATCCTCCTCCTCCGTTGATGAGGATTCCAGCAGAT-3′. The resulting PCR product was ligated into the pDONR221 middle entry Gateway vector, generating a pME-*llgl1*CDS vector. A p3E-noATGmCherry entry vector was generated by PCR-amplifying the mCherry sequence using the following primers: F: 5′- GGGGACAGCTTTCTTGTACAAAGTGGTCGTGAGCAAGGGCGAGGAGGATAACA-3′; R: 5′- GGGGACAACTTTGTATAATAAAGTTGTTTACTTGTACAGCTCGTCCATGCCG-3′. The resulting PCR product was cloned into the pDONRP2R-P3 entry vector (Kwan et al., 2007). *llgl1*CDS was subsequently recombined with the p5E:*myl7-*promoter entry vector (Veerkamp et al., 2013) and the p3E-noATGmCherry entry vector into the pDestTol2pA3 destination vector (Kwan et al., 2007) to generate the *pDestmyl7:llgl1-mCherry* construct. Gateway cloning was performed using the Tol2kit via standard protocols (Kwan et al., 2007). 10 pg of pDestmyl7:llgl1-mCherry was co-injected with 20 pg of *tol2* mRNA into the cell of 1-cell stage wild-type embryos. At 3 dpf healthy embryos displaying myocardial mCherry expression were selected and grown to adulthood. Founder F0 fish were identified by outcrossing and the progeny (F1) displaying the brightest expression were grown to adulthood. The transgenic line was established from F1 adults displaying a Mendelian ratio of transgene transmission. The transgenic line is designated *Tg(myl7:llgl1-mCherry)^sh679^*.

### Immunohistochemistry

Embryos were fixed in 4% PFA with 4% sucrose either overnight at 4°C or for 3 hours at room temperature, and subsequently washed into methanol. After rehydration, embryos were blocked in 0.1% PBS-Triton-X with 10% Goat Serum. Embryos were incubated overnight at 4°C with primary or secondary antibody in blocking solution. The following primary antibodies were used: anti-Caveolin1a (Cell Signalling Technology, D46G3, 1:100); anti-Crb2a (zs-4, ZIRC, 1:50); anti-GFP (Aves labs, GFP-1020, 1:500); anti-Laminin (Sigma, L9393, 1:100). Following removal of the secondary antibody, embryos were dissected, mounted in Vectashield, and imaged on a Nikon A1 confocal at x40 magnification with a 1µm z-slice interval.

### Morpholino oligonucleotide-mediated gene knockdown

Wt1a was depleted using a previously-published splice-blocking *wt1a* MO targeting the first splice site of the *wt1a* gene (sequence: AAAGTAGTTCCTCACCTTGATTCCT) (Perner et al., 2007). The standard control MO targeting human beta-globin intron mutation was used as a negative control (GeneTools, sequence: CCTCTTACCTCAGTTACAATTTATA). All morpholinos were supplied by GeneTools and diluted to a 1 mM stock in MQ water. Working concentrations were as follows: *wt1a* - 200nM, control - 200 nM, Embryos were injected with 1 nL of morpholino solution.

### In situ hybridisation

Embryos were fixed overnight in 4% PFA, washed in PBST and transferred stepwise into 100% MeOH for storage at -20C. mRNA in situ hybridization was carried out as previously described (Noël et al., 2013). Previously published mRNA *in situ* hybridisation probes are as follows: *lamb1a* and *lamc1* (Derrick, Pollitt, et al., 2021), *wt1a* (Bollig et al., 2006), *myl7* (Yelon et al., 1999). Riboprobes were transcribed from linearized template in the presence of DIG-11-UTP or Fluorescein-11-UTP (Roche). To analyse myocardial vs epicardial gene expression, stained embryos were transferred from methanol to a 30% sucrose solution and agitated overnight. The samples were then frozen in a 1cm mold using O.C.T. Embedding Matrix for Frozen Sections (Cat. no. R40020-E, Pyramid Innovation Ltd) on a copper plate cooled with Liquid Nitrogen.

### Quantification of heart rate

Embryos were transferred in batches of 4 (2 siblings, 2 mutants) into E3 and 4.2% tricaine from a 28.5°C incubator to a dissection microscope attached to a high-speed camera (Chameleon3 USB3, FLIR Integrated Imaging Solutions). The heart of each embryo was located and image sequences (.tif) were captured for up to 20 seconds at 150 frames per second using SpinView Software (Spinnaker v. 2.0.0.147). Image sequences were imported into Fiji and a line drawn through the heart, from which a kymograph was generated to visualise periodicity of cardiac contraction. Heart rate was quantified from kymographs. Individual values represent an average heart rate over a 60 second period.

### Live lightsheet imaging

Embryonic zebrafish hearts were imaged live using a Zeiss z.1 lightsheet microscope. The embryos were anaesthetised and heart contractility arrested by incubating the embryos in 8.4% tricaine in E3 at 10°C. Z-stacks encompassing the entire heart were acquired as previously described (Derrick, Sánchez-Posada, et al., 2021).

### Image quantification

Prior to quantification, image files were blinded using an ImageJ Blind_Analysis plugin (modified from the Shuffler macro, v1.0 26/06/08, Christophe Leterrier, Aix-Marseille University, France). Fluorescent signal intensities were measured using ImageJ.

*Looping ratio* was quantified as previously described (Derrick, Pollitt, et al., 2021) by dividing the looped distance between the two cardiac poles by the linear distance.

*Extruding cell number* was quantified by manually inspecting each z-stack and defined as a cell projecting from the apical surface sufficient to perturb the contour of the external ventricular wall.

*Apical Laminin deposition* was quantified by measuring Laminin fluorescence intensity across the apicobasal axis of myocardial cells, based on a previously described method (Gentile et al eLife, 2021). *myl7:LifeActGFP* signal was also quantified to define the apicobasal geometry of each cell, a threshold was set for GFP intensity that defined the apicobasal limits of the cell. This signal was used to coerce all cells to a uniform width with a range of 0 (basal) to 1 (apical), standardising cell size and facilitating relative analysis of Laminin distribution at basal or apical sites irrespective of cell size. Laminin signal intensity was binned to basal (-0.05 to 0.3) and apical regions (0.7 to 1.05) of each cell, allowing comparison across regions and genotypes. Each cell represents an individual experimental unit.

*Junction or apical membrane Crb2a localisation* was quantified by drawing a 5px thick line was drawn across the apical surface of individual cardiomyocytes, from one cell-cell junction to another, using the *myl7:LifeActGFP* signal as a landmark for cell-cell junctions. Similar to the process described for Laminin, cells were coerced to the same geometry of width (0 to 1), and the Crb2 signal was binned to junctional (0 to 0.2 and 0.8 to 1) and apical membrane (0.2 to 0.8) regions of each cell. Each cell represents an individual experimental unit.

*Intranuclear distance* was computed by first performing live light-sheet imaging of hearts carrying the *Tg(-5.1myl7:DsRed2-NLS)^f2^* transgene to generate 3D z-stacks in which myocardial nuclei express dsRed. Nuclei coordinates were generated using the Imaris spot detection tool. Nuclei coordinates were imported into *morphoCell*, where ventricular nuclei were annotated, non-overlapping nuclear clusters comprised of a central-seed cell and four nearest neighbouring cells were assigned, and the Euclidean distance between the central-seed cell and its neighbouring cells in each cluster automatically measured and averaged, then averaged for all ventricular clusters per embryo.

*Statistical analysis* of quantitative data was performed in Graphpad Prism 9 or using the Statix package in R. Data was tested for normality to determine the most appropriate statistical test. Data were considered significant when *p-value*<0.05.

