## Supplemental Data for "Llgl1 mediates timely epicardial emergence and establishment of an apical laminin sheath around the trabeculating cardiac ventricle"

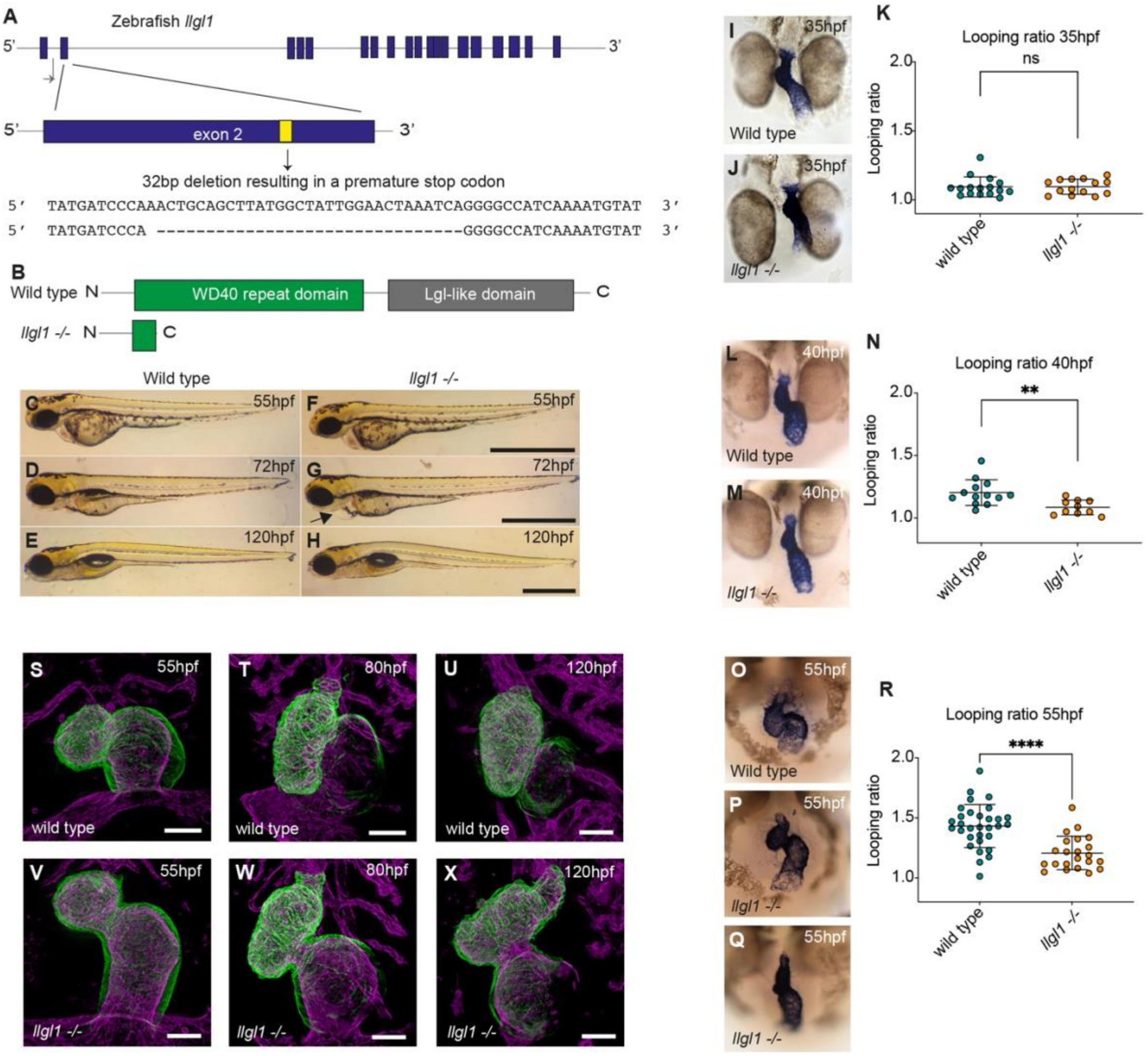


**Figure S1 – Generation of an *llgl1* zebrafish mutant**

A: CRISPR-Cas9-mediated mutagenesis targeting exon2 of zebrafish *llgl1* resulted in a 32bp deletion in exon 2. B: The 32bp *llgl1* deletion results in a premature stop codon generating a truncated protein lacking all functional domains of Llgl1 C-H: Brightfield images of *llgl1* mutants, lateral view, anterior to left. Scale bar = 1mm. *llgl1* mutants exhibit mild cardiac oedema at 72hpf (arrowhead, G) which resolves in most embryos by 5dpf (H). I-R: Analysis of heart looping morphogenesis in wild type, heterozygotes, and *llgl1* mutant embryos between 35hpf and 55hpf using mRNA *in situ* hybridisation analysis of *myl7* expression and quantification of looping ratio (K, N, R, see Materials and Methods for details). At 35hpf, there is no difference in looping ratio between genotypes (I-J: wt n=16; *llgl1* -/- n=14). By 40hpf *llgl1* mutants exhibit a significant reduction in looping ratio compared to wt siblings (L-N: wt n=13; *llgl1* -/- n=10). At 55hpf the looping defect in *llgl1* mutants is visibly more pronounced (O-R: wt n=31; *llgl1* -/- n=22). Heart morphology in *llgl1* mutants at 55hpf is variable, with some mutants exhibiting milder looping disruption (P), while others exhibit severe morphological defects (Q). Kruskal-Wallis test, **** $p<0.0001$, ** $p<0.01$. S-X: live lightsheet microscopy of

*Tg(myl7:LifeAct-GFP);Tg(fli1a:AC-TagRFP)* double transgenic wild-type and *llgl1* mutant embryos, highlighting the myocardium (green) and endocardium (magenta). *llgl1* mutants exhibit ongoing defects in heart morphogenesis from 55hpf (V), including smaller chambers and reduced looping morphology at 80hpf (W) and 120hpf (X), when compared to wild-type siblings (S, T, U). Scale bar = 50μm

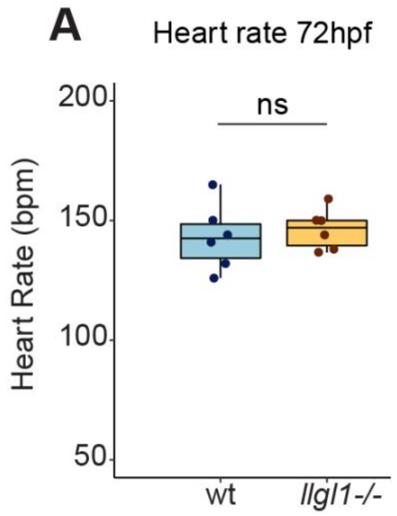


**Figure S2 – Heart rate is unaffected in *llgl1* mutants.**

A: Quantification of heart rate in wild-type siblings (n=6) and *llgl1* mutants (n=6) at 72hpf. No significant difference in heart rate is observed in *llgl1* mutants. T-test, ns = not significant.

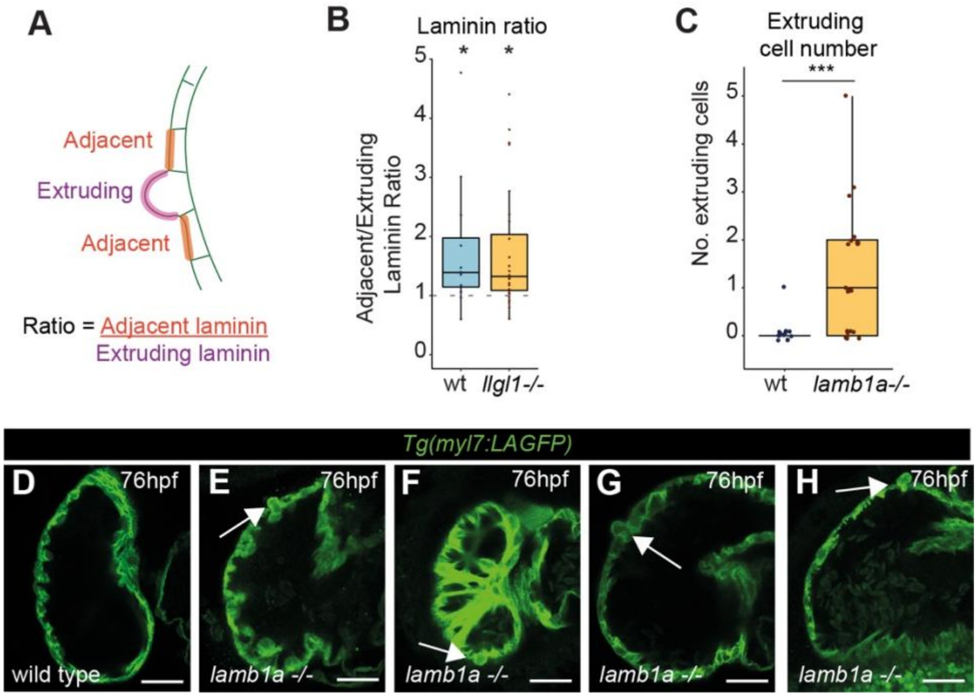


**Figure S3 - *lamb1a* mutants exhibit apical ventricular CM extrusion.**

A: Quantification of adjacent cell/extruding cell Laminin ratio in both wild-type sibling and *llgl1* mutant embryos. A value of greater than 1 indicates lower apical Laminin in extruding cells than adjacent cells. 1-sided T-test, * $p<0.05$. B: Schematic depicting quantification of adjacent/extruding Laminin ratio. C: Quantification of extruding cell number at 80hpf in wild-type siblings (n-13) and *llgl1* mutant embryos (n=22). T-test, *** $p<0.001$. D-H: Confocal z-slices through the ventricle of *Tg(myl7:LifeAct-GFP)* transgenic embryos visualising the myocardial wall in wild-type (D) and *lamb1a* mutant embryos (E-H) at 76hpf. Scale bar = 25µm. Cardiomyocytes can be seen extruding apically from the ventricular wall in *lamb1a* mutants (arrowheads E-H).

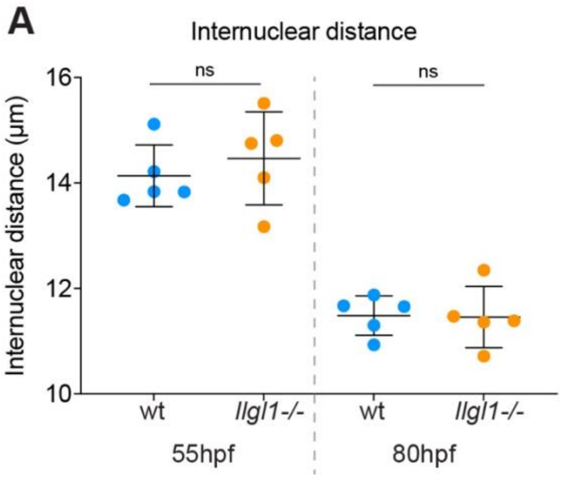


**Figure S4 - Ventricular cardiomyocyte size is unaffected in *llgl1* mutants.**

A: Quantification of ventricular cardiomyocyte internuclear distance as a proxy for cardiomyocyte size wild-type embryos and *llgl1* mutants at 55hpf and 80hpf (n=5 for each stage/genotype). Each point represents the average internuclear distance of the ventricular cells in an individual embryo. Cell size is comparable between wild-type embryos and *llgl1* mutants at both stages. T-test, ns = non significant.

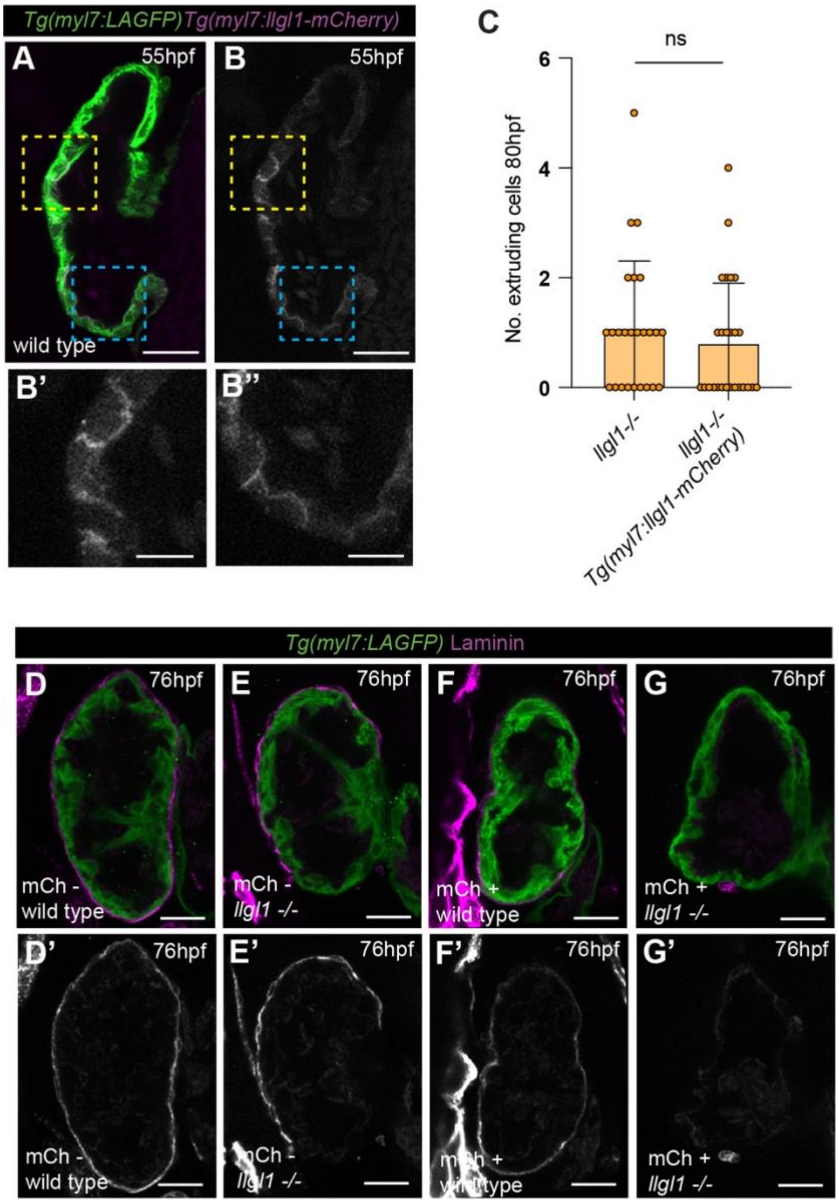


**Figure S5 - Myocardial re-expression of *llgl1* in *llgl1* mutant embryos does not rescue cell extrusion or epicardial defects.**

A-B: Confocal z-slices of the ventricle of *Tg(myl7:LifeAct-GFP), Tg(myl7:llgl1-mCherry* transgenic embryos visualising the myocardium (green) and mCherry-tagged *llgl1* expressing myocardial cells (magenta A, B) at 55hpf. Scale bar = 25µm. B' and B'' show higher magnifications of the yellow and blue dashed boxes, respectively, shown in panels A,B, scale bar = 10µm. B,B'' Llgl1-mCherry is enriched at the basement membrane of cardiomyocytes. C: Quantification of extruding cell number in *Tg(myl7:llgl1-mCherry)*-positive *llgl1* mutant embryos (n=26) and transgene-negative *llgl1* mutant embryos (n=25) at 80hpf. No significant difference in the number of extruding cells is observed between *Tg(myl7:llgl1-mCherry)*-positive or -negative *llgl1* mutants (Kruskal-Wallis). D-G: Confocal z-slices of the ventricle of *Tg(myl7:LifeAct-GFP)* transgenic embryos visualising the myocardium (green) stained with an anti-Cav1a antibody (magenta) in *Tg(myl7:;llgl1-mCherry)*-positive wild-type (D) and *llgl1* mutant embryos (E), and *Tg(myl7:llgl1-mCherry)*-negative wild-type siblings (F) and *llgl1* mutant embryos (G) at 76hpf. Scale bar = 25µm. Both *myl7:llgl1-mCherry*-positive wild-type (n=7/9) and *myl7:llgl1-mCherry*-negative wild-type embryos (n=11/14) have full epicardial coverage. As expected, a subset of *myl7:llgl1-mCherry*-negative *llgl1* mutant embryos exhibit only partial epicardial coverage (n=10/16). *myl7:llgl1-mCherry*-positive *llgl1* mutant embryos, in which Llgl1 is returned to the myocardium, exhibit similar numbers of embryos with epicardial defects (n=14/16).
